## Supplementary Figures for "Targeted sensors for glutamatergic neurotransmission"

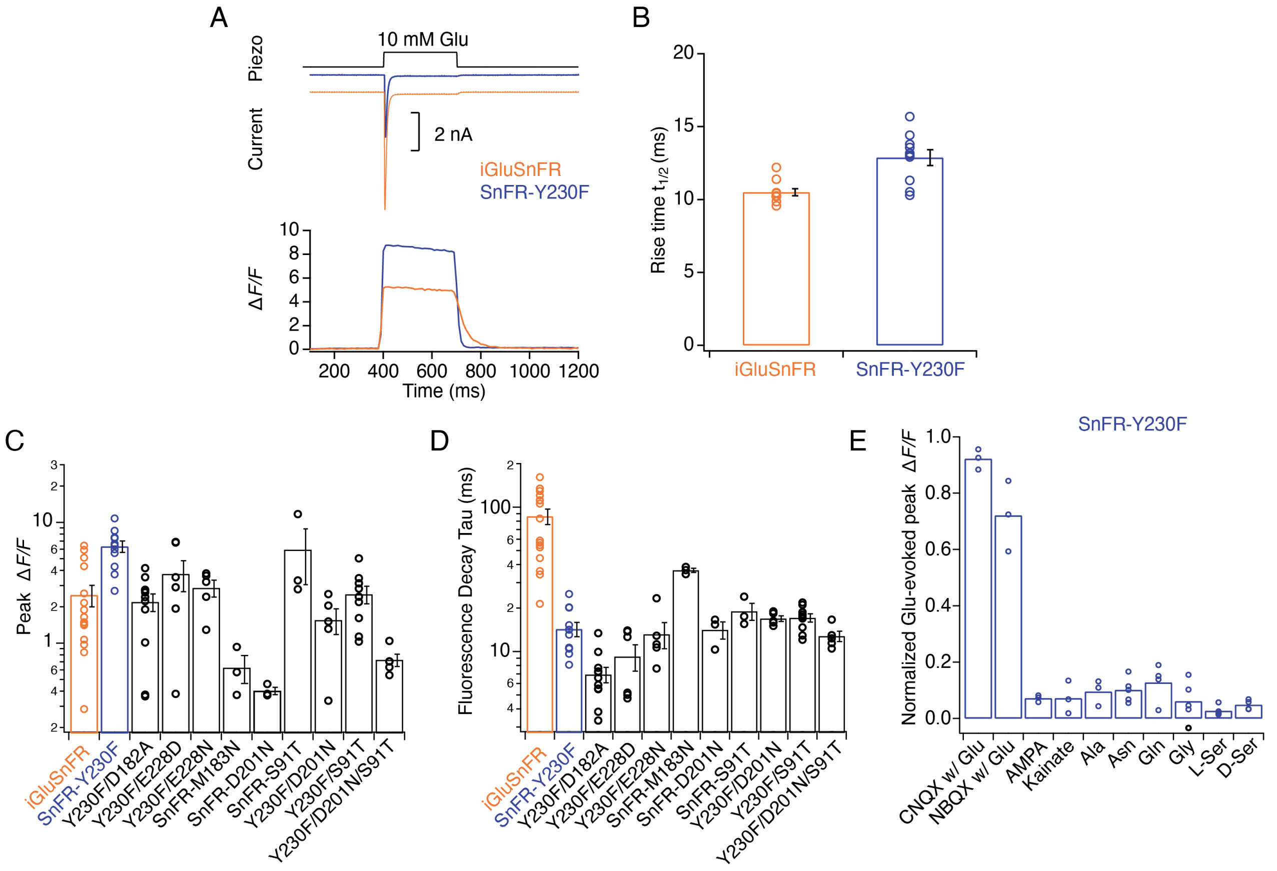


**Figure S1**. **Kinetic properties and selectivity of iGluSnFR variants in HEK cells.** (A) Patch clamp fluorometry of iGluSnFR (orange traces) and the SnFR-Y230F variant (blue traces) in whole lifted cells revealed a stronger fluorescence response and faster off kinetics for Y230F than the iGluSnFR parent construct. Representative current traces show the independent response to 10 mM glutamate from co-transfected GluA2 AMPA receptors, indicating the fast solution exchange (~5 ms). (B) Rise time to peak fluorescent response of iGluSnFR for 10 mM glutamate application ($n$ = 10) and Y230F ($n$ = 10). (C)-(D) Statistics of peak fluorescent response and decay time constant at the end of the glutamate pulse of iGluSnFR and mutants. (E) Statistics of normalised effects of GluA2 inhibitors, agonists and amino acids on SnFR-Y230F variant in patch clamp fluorometry, indicating it retains the excellent specificity of the parent construct.

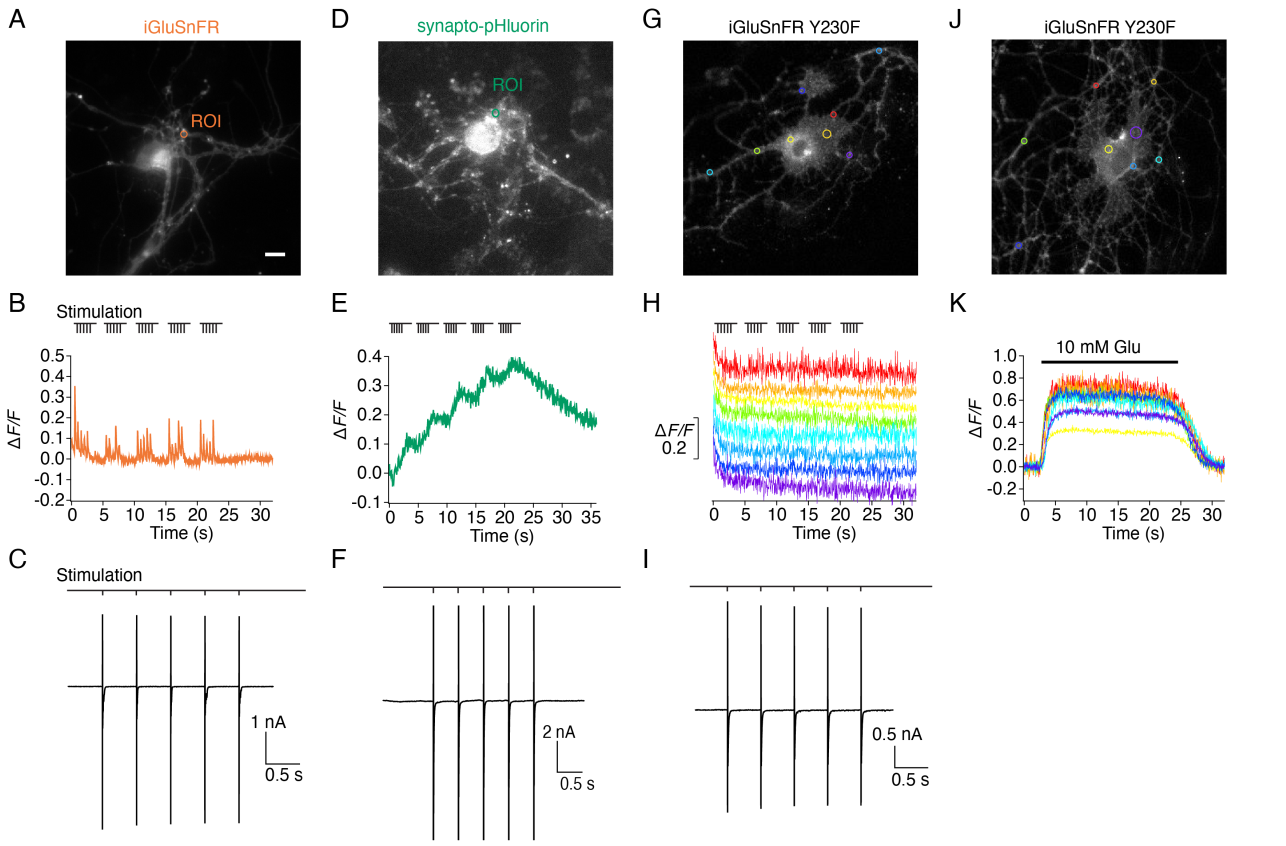


**Figure S2. Evoked currents and fluorescence signals of iGluSnFR, SnFR-Y230F variant and synapto-pHluorin on autaptic neurons.** (A) Representative fluorescence micrograph (scale bar, 5 μm) for an autaptic neuron expressing iGluSnFR with a region of interest (ROI) marked. (B) Fluorescence time series from the region of interest recorded during a patch clamp experiment. (C) Excitatory post-synaptic currents were evoked by escaping action potentials (also drawn). Stimulation of 5 pulses at 2 Hz shows robust iGluSnFR responses. (D-F) As for panels (A-C) but for an autaptic neuron expressing synapto-pHluorin, showing an accumulating fluorescence increase from the ROI marked with a green circle in (D) that returned much more slowly to the baseline. (G-I) As for panels (A-C) but for an autaptic neuron expressing the SnFR Y230F mutant. Evoked currents of Y230F variant were similar to those seen for iGluSnFR and synapto-pHluorin, but there were no fluorescence signals at any region of interest during stimulation with the same protocol. 8 colour-coded time series are shown, corresponding to the coloured circles in panel G. (J & K) Direct application of 10 mM glutamate close to the neuron gave robust fluorescence increases at multiple sites, indicating that the SnFR Y230F mutant was membrane expressed and was functional.

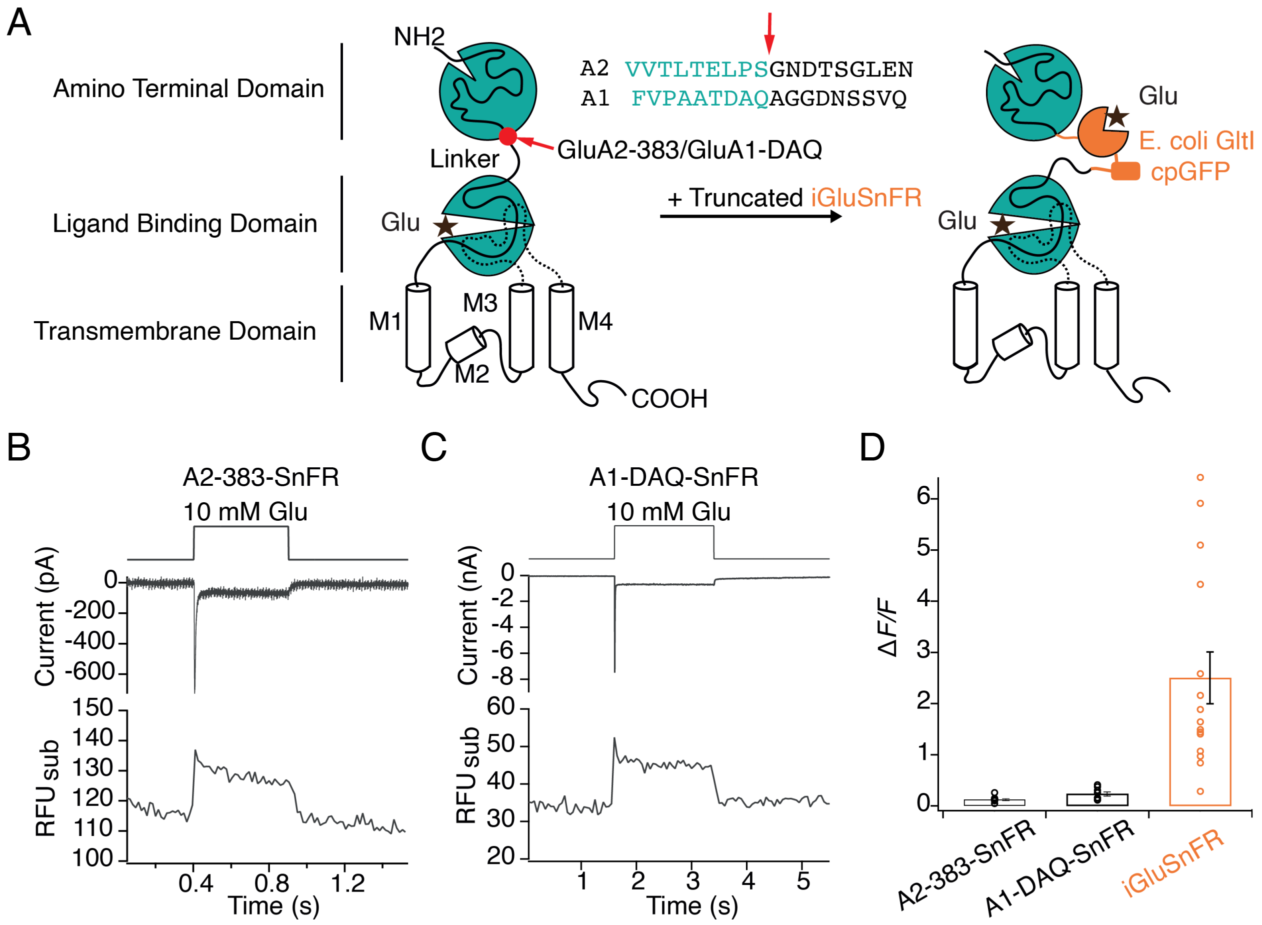


**Figure S3**. **Poor optical performance of truncated SnFR insertions to chimeric AMPA receptors.** (A) Engineering of GluA - iGluSnFR chimeras. Truncated iGluSnFR (GltI and cpGFP), flanked by short linker sequences was inserted at 383 and DAQ sites at the beginning of the linker between ATD and LBD domains of GluA2 and GluA1. Insertion points in the amino acid sequences of GluA2 and GluA1 receptors are indicated in red. (B)-(C) Representative whole cell current and sub-region fluorescent traces for GluA2-383-SnFR and GluA1-DAQ-SnFR, showing normal current responses but very small fluorescence changes upon glutamate (10 mM) perfusion. (D) Bar graph showing that fluorescence responses to glutamate for GluA2/A1 SnFR chimeras are far weaker than those of iGluSnFR in patch clamp fluorometry.

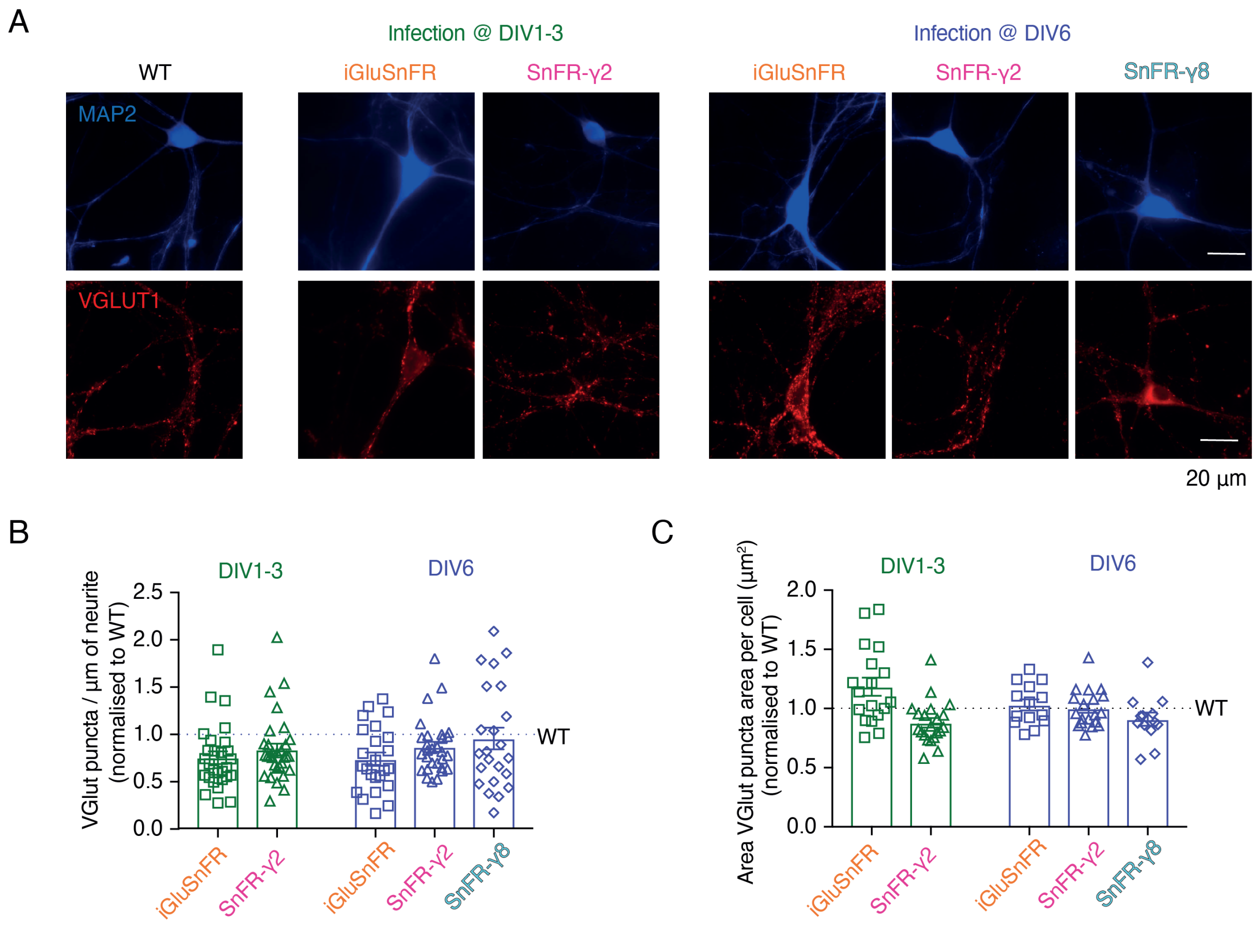


**Figure S4. Neither SnFR-γ2 nor SnFR-γ8 influence synapse density or size in hippocampal neuron cultures.** (A) Representative images of synapse density in control (WT), iGluSnFR, SnFR-γ2 and SnFR-γ8 expressed in bulk hippocampal neuron cultures infected at DIV1-3 (middle panel) or DIV 6 (right panel). Synapse density was assessed by double immunofluorescence labelling for MAP2 (blue, top images) and VGLUT1 (red, bottom images). Scale bar, 20 µm. (B & C) Quantification of synapse density per µm of neurite (B) and area of VGLUT puncta per cell (C) in neurons infected with iGluSnFR and SnFR-γ2 at DIV1-3 ($n$ = 18 and 21 respectively, green bars) and neurons infected with iGluSnFR, SnFR-γ2 and SnFR-γ8 at DIV6 ($n$ = 14, 19 and 13 respectively, purple bars) normalised to WT (n= 16). Data shown are individual points and means with standard deviation of the mean. Three independent neuronal cultures were assessed, and the statistical test of no difference was the Mann-Whitney nonparametric test.

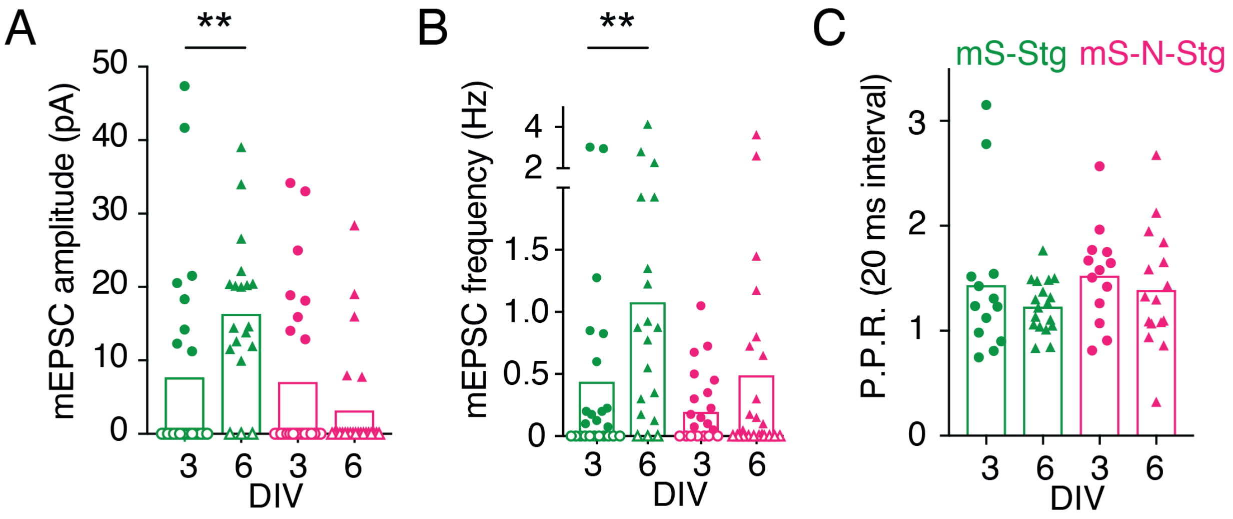


**Figure S5**. **Electrophysiological parameters of autapses overexpressing Stargazin constructs** Miniature EPSC amplitude (A), mEPSC frequency (B), and paired-pulse ratio (P.P.R., C) for neurons expressing mScarlet-Stg (green) or mScarlet-NETO-Stg (magenta) that were infected at either 3 (circles) or 6 (triangles) days in vitro (DIV). For (A) and (B), three independent neuronal cultures were examined (from left to right, $n$ = 25, 24, 20 and 26 cells, respectively), with recording and analysis performed blind to the construct used. Open symbols correspond to zero values from excluded cells (see Methods for criteria). For (C), across the four conditions (mS-Stg, DIV 3 and DIV 6 and and mS-N-Stg, DIV 3 and DIV 6), 10/23, 2/20, 11/24 and 3/24 cells were non-responsive. Data shown are individual points, bars represent means. Probability of no difference is from the Mann-Whitney non-parametric test (** $p < 0.01$).

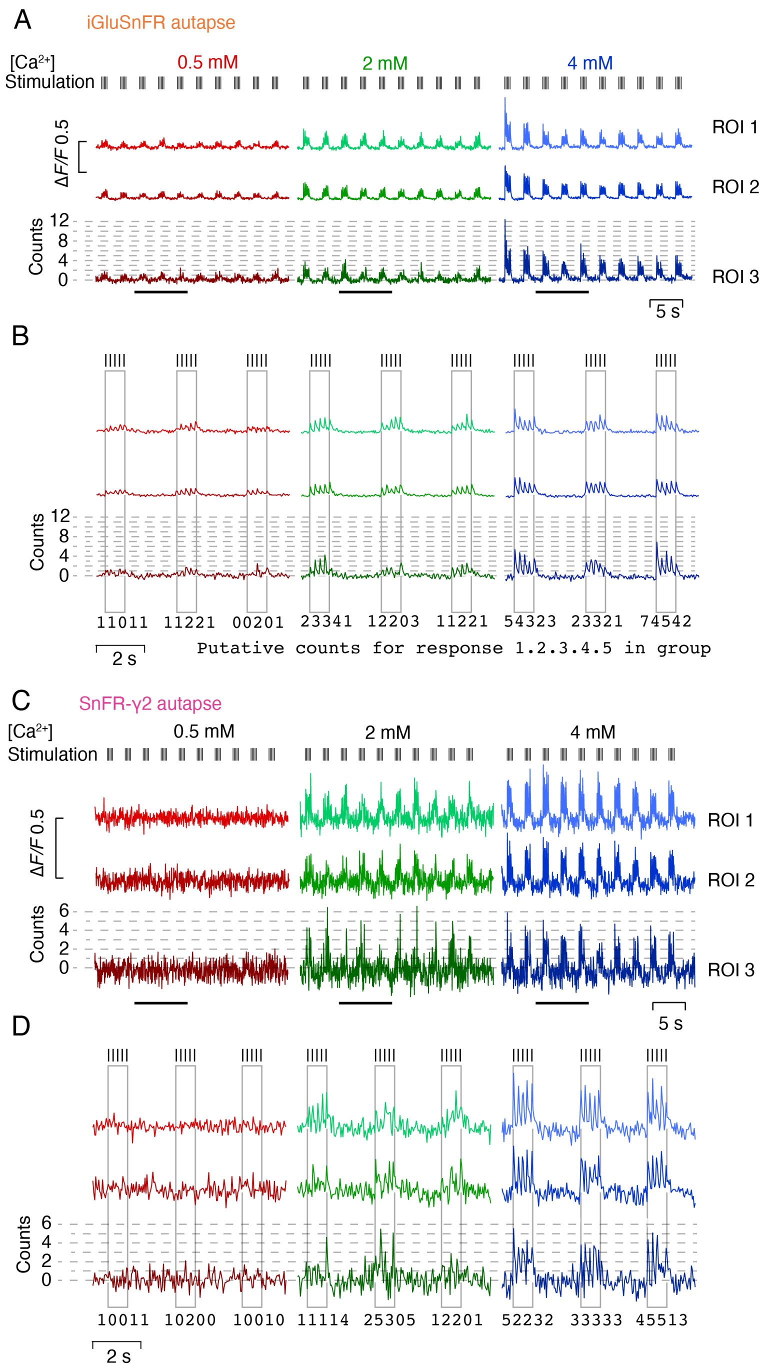


**Figure S6 Quantal nature of indicator responses** (A) Fluorescence time series at 3 calcium concentrations from 3 ROI in an autapse expressing iGluSnFR. Quantal analysis for ROI 3 is shown in Figure 8. Note the development of steady-state responses and mild rundown. (B) Zoom in to 3 groups of responses indicated by bars in panel (A). For each stimulation, a putative quantal count is given. We did not count quanta in this way, rather binning peak amplitudes into histograms. Note that the agreement of the peaks to the dashed lines is imperfect. (C) As for panel A but for a cell expressing SnFR-γ2. Responses have smaller signal to noise ratio but are more consistent, showing no rundown and better quantal nature. Again, ROI3 is analysed in Figure 8. (D) As for panel B but for the same cell as in panel (C).

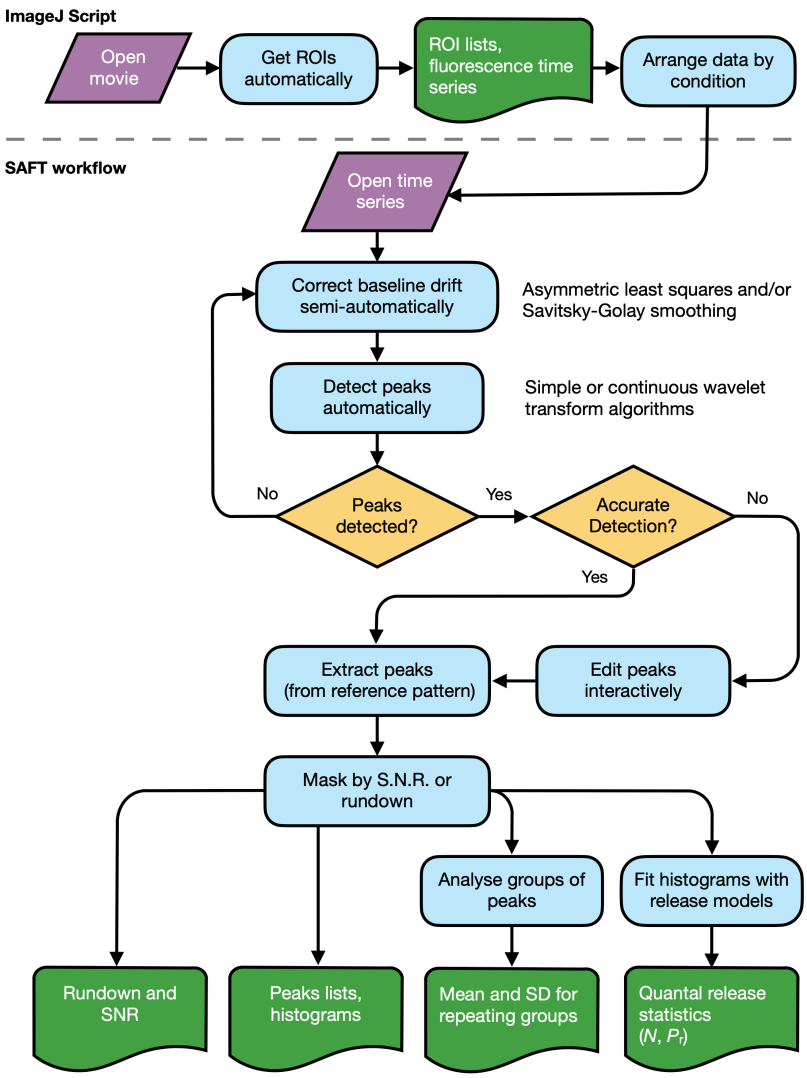


**Figure S7 Workflow for peak response extraction and analysis of fluorescent report of glutamate release from synapses** A script in imageJ is used to detect regions of interest with changes in fluorescence over the course of movies. This determination exploits information about the number of expected responses if it is available. The data is organised into spreadsheets of time series, with each sheet corresponding to a different condition in the same experiment. These data are loaded into the PYTHON application SAFT. Baseline drift and peak detection are automated but supervised and interactive for the user. SAFT generates an "mean" trace to enable precise identification of the expected peak locations. These peak locations are used to extract the responses from an arbitrary number of ROIs. The output can be masked according to rundown or low signal to noise, and the listed peak responses can be analysed in groups or individually to determine quantal parameters for each putative synaptic site.

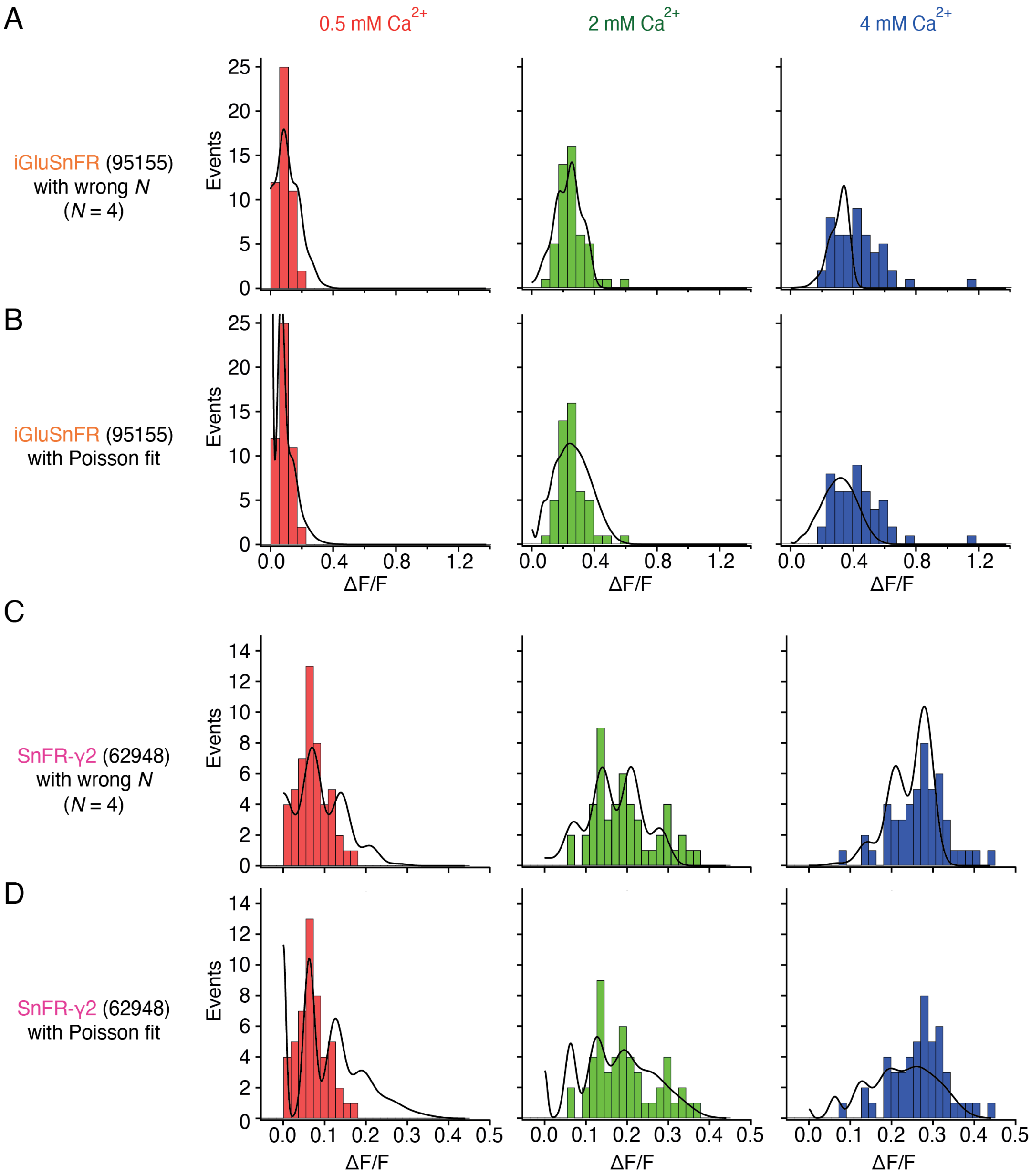


**Figure S8 Examples of imperfect fits to peak response histograms** (A) For a binomial model, fixing the number of releasable vesicles (*N*) too small fails to describe the full width of the distribution at 4 mM $Ca^{2+}$ . At 4 mM $Ca^{2+}$, the release probability was erroneously determined to be 89% in this fit, because the full width of the data were not described. (B) Poisson fit to the same data as in panel (A) is featureless, due to narrow optimised quantal size. Also, the number of failures ($\Delta F/F \approx 0$) is overestimated at 0.5 mM $Ca^{2+}$ (because release rate was optimised to be 0.96 for this condition). (C) A fit to SnFR-γ2 data with *N* too small overestimates $P_R$ in all conditions (29%, 60%, 85% in 0.5, 2 and 4 mM $Ca^{2+}$ respectively), without describing the full range of observed responses. (D) Poisson fit to the same data as in panel (C) erroneously overestimates failures at 0.5 mM and 2 mM $Ca^{2+}$, predicts multi vesicular release at 0.5 mM and underestimates larger events at 4 mM $Ca^{2+}$.
