## Supplementary Code - ImageJ Macro for "Targeted sensors for glutamatergic neurotransmission": SnFR_ROI_Finder.ijm

```
//script to find ROIs in a Stack with clear defined peaks
// you need to have two plug-ins "Find Peaks" and "Read and Write to excel"
// get them by clicking on help->update, than "manage update sites" tick there
// BAR and Result to Excel
//Open a stack to be analyzed and run the script :)
// Clear everything to make sure no errors occur
run("Clear Results");
nRois=roiManager("count");
if(nRois>0){
roiManager("reset");
}
//Remember Stack Title and get dimensions, measure mean overall intensity
Dialog.create("Setting your parameters");
Dialog.addNumber("Lenght of the sides of the ROI", 7);
Dialog.addNumber("Intensity increase upon an event (e.g. 10%=0.1)", 0.1);
Dialog.addNumber("Signal to Noise Ratio", 3);
Dialog.addNumber("Minimal number of expected Peaks", 10);
Dialog.show();
dir=getDirectory("Choose where your Rois shall be stored");
SizeofRoi=Dialog.getNumber();
IncreaseFactor=Dialog.getNumber();
SNFactor=Dialog.getNumber();
ExpectedPeaks=Dialog.getNumber();
run("Subtract Background...", "rolling=50 stack");
T=getTitle();
getDimensions(width, height, channels, slices, frames);
Width= width/SizeofRoi;
Height= height/SizeofRoi;
selectWindow(T);
run("Select All");
run("Measure");
MeanofAll= getResult("Mean",0);
run("Clear Results");
//Enter x,y run
u=0;
xRoi=0;
yRoi=0;
Roicount=0;
t=0;
while(t<Height){
	
while(u<Width){

selectWindow(T);
makeRectangle(xRoi, yRoi, SizeofRoi, SizeofRoi);
roiManager("add");
run("Measure");
MeanofRoi= getResult("Mean", 0);
run("Clear Results");
//test if mean intensity in current Roi is lower than mean overall intensity
//go to next iteration if so
if(MeanofRoi<=MeanofAll){
roiManager("select", Roicount);
roiManager("delete");
xRoi=xRoi+SizeofRoi;
u=u+1;
}
else{

selectWindow(T);
roiManager("select", Roicount);
roiManager("Multi Measure");
selectWindow("Results");
Table.sort("Mean1");
n=0;
o=nResults-1;
Max=0;
Min=0;
Maximum=0;
Minimum=0;
meanMax=0;
meanMin=0;
Amplitude=0;
while(n<ExpectedPeaks){
	Max= getResult("Mean1", o);
	n= n+1;
	Maximum= Maximum+Max;
	o=o-1;
}
v=0;
while(v<30){
	Min= getResult("Mean1", v);	
	v=v+1;
	Minimum= Minimum+Min;
}


meanMax= (Maximum)/ExpectedPeaks;
meanMin= (Minimum)/30;
Amplitude= (meanMax-meanMin)*0.5;
MinimumIncrease= meanMin*IncreaseFactor;


if(Amplitude<MinimumIncrease){
roiManager("select", Roicount);
roiManager("delete");
xRoi=xRoi+SizeofRoi;
u=u+1;
close("Results");


}
else{
//clean up
close("Resutls");
//determine bin with highest counts in intensity to not fall into a mean value that is artificially high due to good signals
selectWindow(T);
roiManager("select", Roicount);
roiManager("Multi Measure");
run("Distribution...", "parameter=Mean1 or=16 and=0-0");
Plot.getValues(bin, count);
Array.show("Results",bin, count);
selectWindow("Results");
Table.sort("count");
close("Mean1 Distribution");
RealMean=getResult("bin", 15);
Amplitude=meanMax-RealMean;


//do not use Rois with an peak amplitude that is smaller than 10 % of mean intensity
Mini=RealMean*IncreaseFactor;
if(Amplitude<Mini){
roiManager("select", Roicount);
roiManager("delete");
xRoi=xRoi+SizeofRoi;
u=u+1;
close("Results");
}

else{
MeanToMin=(RealMean-meanMin)*SNFactor;
if(Amplitude<MeanToMin){
roiManager("select", Roicount);
roiManager("delete");
xRoi=xRoi+SizeofRoi;
u=u+1;
close("Results");	
}

else{
roiManager("select", Roicount);
roiManager("Multi Measure");
roiManager("select", Roicount);
Plot.create("Plot of Results", "x", "Mean1");
Plot.add("Line", Table.getColumn("Mean1", "Results"));
Plot.setStyle(0, "blue,#A0A0FF,1.0,Line");
Plot.show();
SearchAmplitude=Amplitude*0.6;
run("Find Peaks", "min._peak_amplitude=SearchAmplitude min._peak_distance=0 min._value=[] max._value=[] list");
IJ.renameResults("Plot Values", "Results");
MinimumNrPeaks=ExpectedPeaks/3;
Test=0;
Test=getResult("Y1", MinimumNrPeaks);
Test2=0;
Test2=getResult("Y1",50);
if(isNaN(Test)){
	roiManager("select", Roicount);
roiManager("delete");
xRoi=xRoi+SizeofRoi;
u=u+1;
close("Results");
close("Peaks in Plot of Results");
close("Plot of Results");
}

else{

if(isNaN(Test2)){
close("Peaks in Plot of Results");
roiManager("select", Roicount);
rName= Roi.getName();
selectWindow("Plot of Results");
Plot.freeze();
rename(rName);
Roicount=Roicount+1;
xRoi=xRoi+SizeofRoi;
u=u+1;	
close("Results");
run("Tile");
}
else{
	roiManager("select", Roicount);
roiManager("delete");
xRoi=xRoi+SizeofRoi;
u=u+1;
close("Results");
close("Peaks in Plot of Results");
close("Plot of Results");			
}
						}// end of else if(Amplitude<MeantoMin)
					}//end of else 	(isNaN(Test))
			}//end of else (Test is a real Number, so Image contains sufficient peaks
}
		}//end of else (MeanofRoi<=MeanofALL)
	
	} //end of u<width while loop
u=0;
t=t+1;
xRoi=0;
yRoi=yRoi+SizeofRoi;
}//end of t<height while loop
//User should have a look on the tiled traces and manually delete bad ones
title="Decision Time";
message="Inspect tiled traces for outliers, if you found one, check the name of the plot to find it in the Roi Manager and delete it there. Click OK when finished";
close("Results");
waitForUser(title, message);
//run through all rois sequentially and multi measure over stack, write each measurement to excel
nRoi= roiManager("count");
p=0;
run("Set Measurements...", "mean redirect=None decimal=3");
selectWindow(T);
for(p=0; p<nRoi; p++){
roiManager("Select", p);
roiManager("Multi Measure");
run("Read and Write Excel", "no_count_column");
}
roiManager("Show All without labels");
selectWindow(T);
saveAs("Tiff", dir+"/overlay"+T+".tiff");
run("Close All");
roiManager("Deselect");
roiManager("Save", dir+"/RoiSet"+T+".zip");
roiManager("Deselect");
roiManager("delete");
```
